# Ascitic fluid protects against ferroptosis and enables the peritoneal spread of ovarian cancer

**DOI:** 10.1101/2024.11.23.624998

**Authors:** Yasaman Setayeshpour, Ssu-Yu Chen, Divya L. Dayanidhi, Yunji Lee, Chiara Federico, Juan J. Aristizabal-Henao, Jianli Wu, Chao-Chieh Lin, Alexander A. Mestre, Michael A. Kiebish, Andrew Berchuck, David S. Hsu, Zhiqing Huang, Susan K. Murphy, Jen-Tsan Chi

## Abstract

One of the most common sites of metastasis in ovarian cancer (OVCA) is the peritoneum. Often, this spread is accompanied by the accumulation of a fluid called ascites in the peritoneal cavity. Despite its common occurrence in metastatic OVCA patients, ascites and its influence on the peritoneal spread of OVCA are poorly understood. Interestingly, OVCA cells are vulnerable to ferroptosis, a type of cell death caused by lipid peroxidation. Hence, how these ferroptosis-sensitive OVCA cells persist in their spread to the peritoneum remains unknown. Here, we show that ascites robustly protects OVCA cells and patient-derived organoids against ferroptosis and enhances the peritoneal spread of OVCA cells in mice. Mechanistically, ascites downregulates the mitochondrial enzyme, 3-hydroxy-3-methylglutaryl-CoA synthase 2 (*HMGCS2*), which contributes to an increase in lipid droplets. Additionally, upon ferroptosis induction, ascites represses the upregulation of the transferrin receptor, *TFRC*, thereby decreasing cellular labile iron levels. Furthermore, we show that lipid-lowering fibrates reverse cellular changes induced by ascites, and they attenuate the peritoneal spread of OVCA cells in mice. Our findings implicate the importance of ascites in ferroptosis protection and the peritoneal spread of OVCA, and they suggest that targeting the ferroptosis protection by ascites may present a novel therapeutic approach to limit OVCA metastasis.

## Introduction

Metastasis significantly contributes to the fatal progression of high-grade ovarian cancer (OVCA) (1, 2). OVCA metastasis typically involves the spread of cancer cells from the primary tumors, either in the ovaries or the fallopian tubes, to other parts of the abdominal cavity, such as the omentum (3, 4), the peritoneum (5), and lymph nodes (6). Advanced high-grade OVCA is often accompanied by increased vascular permeability and fluid leakage (1), leading to an abnormal accumulation of the ascites fluid in the peritoneal cavity (1, 7). Although ascites is observed in other diseases, such as liver cirrhosis, it is most frequently observed with metastatic OVCA (1, 7). Due to the enrichment of cellular and acellular factors, the ascitic fluid is reported to harbor a growth-promoting and immune-evading environment for cancer cells (8, 9) and is thought to serve as a medium for cancer cell dissemination and tumor progression and metastasis (1). However, much remains unknown about how ascites affects the process of peritoneal spread of OVCA.

The biological processes in the metastatic cascade for OVCA cells are harsh, often rendering them sensitive to various stressors (5, 9–12). Specifically, we and other groups have reported that detached and metastasizing OVCA cells are especially vulnerable to ferroptosis (13–18), a form of cell death characterized by iron dependency and an irreversible accumulation of lipid peroxides (19). Lipid peroxidation can occur due to ROS (reactive oxygen species) generation, which are natural byproducts of several cellular metabolism pathways and are often catalyzed by labile iron (20). Usually, cellular defense mechanisms are in place to prevent or repair lipid peroxidation, thereby preventing ferroptosis (21, 22). For example, the xCT transporter imports cystine, which leads to the generation of the antioxidant glutathione (GSH) as a cofactor of glutathione peroxidase 4 (GPX4), which repairs lipid peroxidation (21, 22). Concurrently, GPX4-independent ferroptosis protection mechanisms include the ferroptosis suppressor protein 1 (FSP1) and dihydroorotate dehydrogenase (DHODH), which generate ubiquinol to protect against ferroptosis on the plasma membrane and inner mitochondrial membrane, respectively (22). Finally, lipid droplets can protect against ferroptosis by serving as protective sinks (23–26). Nevertheless, ferroptosis ensues if lipid peroxidation accumulation outweighs the cellular neutralizing mechanisms or when these mechanisms are inhibited by various ferroptosis inducers (20). Previously, we have found that OVCA is particularly sensitive to ferroptosis (14). In addition, several metastasis-associated processes promote ferroptosis, such as epithelial-mesenchymal transition (EMT) (27), loss of cell-cell contact (28), the YAP1/TAZ activation (14, 28), and the detachment of tumor cells from adjacent cells and the ECM (29).

This vulnerability of metastasizing OVCA cells to ferroptosis is intriguing, as it confounds how these ferroptosis-sensitive OVCA cells overcome ferroptosis-promoting processes to establish metastasis. To investigate this intriguing question, we wanted to examine the potential role of ascites in ferroptosis sensitivity and the peritoneal spread of OVCA. Nothing is currently known about how ascites may influence OVCA cells’ ferroptosis. For metastasizing melanoma cells, the lymph significantly impacts ferroptosis sensitivity, which may contribute to preferential lymphatic metastasis (30). Given the common occurrence of ascites with peritoneal metastasis, ascites may be crucial for the peritoneal spread of OVCA.

Here, we report that OVCA ascites robustly protects OVCA cells and OVCA patient-derived organoids against ferroptosis, and it enhances the peritoneal spread in murine models. Mechanistically, ascites protects against ferroptosis by increasing lipid droplets via repression of the mitochondrial enzyme, 3-hydroxy-3-methylglutaryl-CoA synthase 2 (*HMGCS2*) and a subsequent decrease in fatty acid oxidation. Furthermore, ascites represses the ferroptosis-induced membrane upregulation of the transferrin receptor (TFRC) and labile iron. We show that these effects of ascites can be attenuated using lipid-lowering fibrates, namely bezafibrate, and we validate our findings in OVCA patient-derived organoids and murine models.

## Results

### Ascites protects ovarian cancer cells from ferroptosis

To investigate whether ascites collected from OVCA patients influence ferroptosis, we added ascitic fluid collected from three different patients with metastatic OVCA to CAOV3 and TYKNU (high-grade serous), TOV21G (clear cells), and TOV112D (endometroid) cells. Ferroptosis was triggered in these cell lines by ferroptosis-inducing agents without or with ascites, and the effects of ascites on ferroptosis were determined via cell viability. As little as 2% of ascites robustly protected these cell lines against ferroptosis induced both by the xCT inhibitor, erastin (**Fig. 1A**), and RSL3, an inhibitor of GPX4 (**Fig. 1B**) (21). This ferroptosis protection by ascites was reproduced with additional ferroptosis inducers that have distinct chemical structures, including Imidazole Ketone Erastin (IKE) and JKE-1674 (**Supplementary Fig. 1A-B**). We also genetically triggered ferroptosis by knocking down *GPX4* and saw that all three tested ascites samples rescued CAOV3 cells from ferroptosis (**Supplementary Fig. 1C**). Furthermore, adding similar levels of fetal bovine serum (FBS) in the media showed no protective effects against ferroptosis, indicating that the rescue effect is not due to changing media composition (**Supplementary Fig. 1E**). To determine the specificity of ascites protection, we tested ascites’ impact on cell viability of CAOV3 cells when treated with non-ferroptosis cell death inducers (e.g., staurosporine, actinomycin D, and rapamycin) and did not observe a robust protection by ascites (**Supplementary Fig. 1F-H**). These results rule out a general protection of ascites against all drug-induced cell death. We also measured the effects of ascites on various ferroptosis biomarkers. While RSL3 and erastin significantly increased lipid peroxidation, this increase was abolished by ascites (**Fig. 1C-D**). Furthermore, the induction of *CHAC1* and *SLC7A11* mRNA during ferroptosis was significantly decreased by ascites co-treatment (**Supplementary Fig. 1I-J**).

**Fig. 1.**
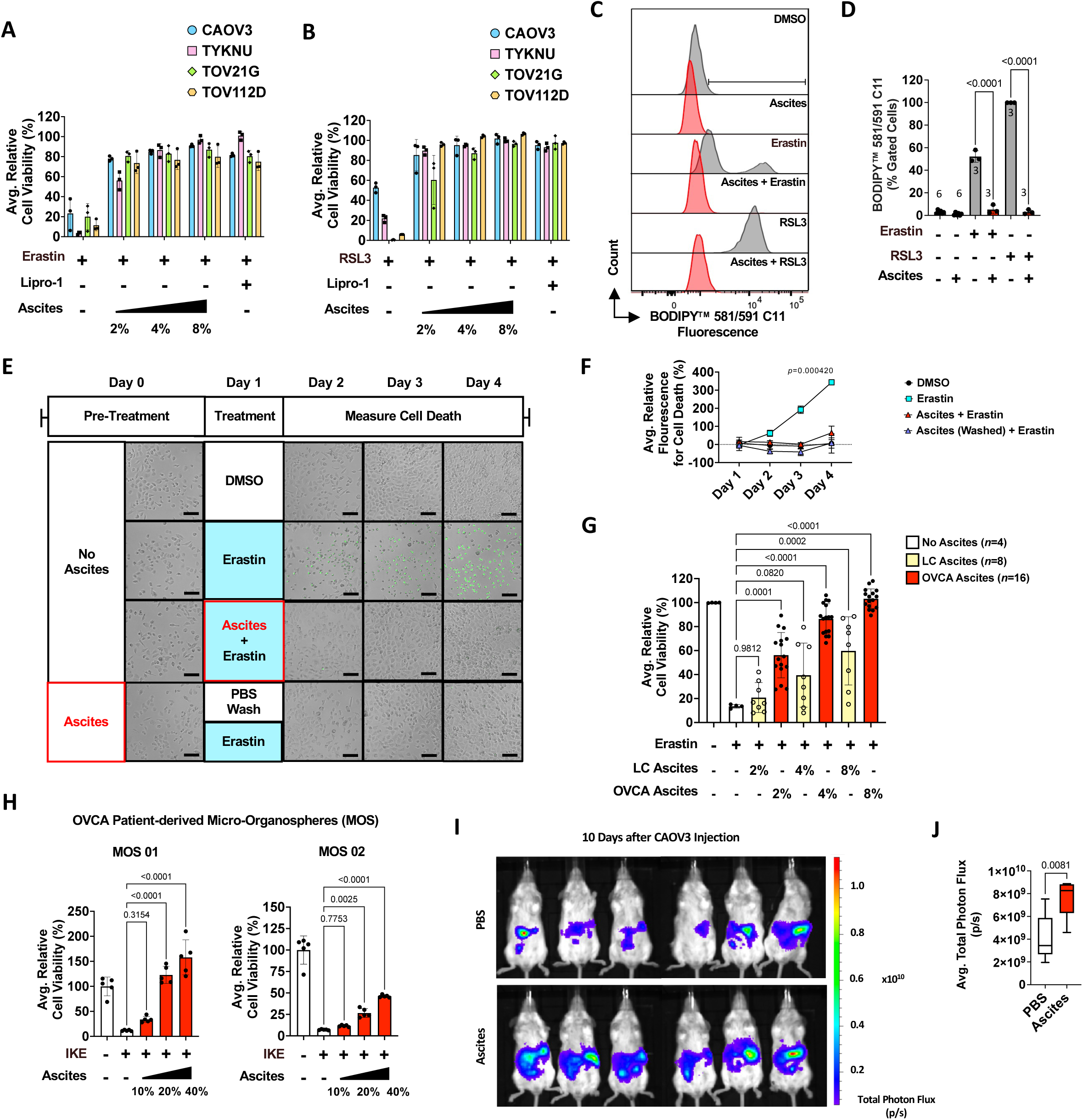
Ascites protects OVCA from ferroptosis. **A-B,** CAOV3, TYKNU, TOV21G, and TOV112D cells were treated with (**A**) 10µM erastin or (**B**) 250nM RSL3 in the presence or absence of 2-8% ascites from three OVCA patients. Liproxtasin-1 (Lipro-1) (2µM) was used as a control (*n*=3/cell line, 24 hours). **C-D**, cells were treated with 5µM erastin for 20 hours or 2µM RSL3 for 2 hours in the presence or absence of 2% ascites. Lipid peroxidation was measured via flow cytometry analysis of BODIPY^TM^ 581/591 C11 staining (*n* provided in panel). **E-F**, cells were either pre-treated with 8% OVCA ascites (Day 0, 16 hours) or left untreated and subsequently treated with 10µM erastin and 8% OVCA ascites. Cell death was visualized (**E**) and measured (**F**) via the CellTox^TM^ Green assay (Days 2-4) in which green fluorescence indicates cell death (*n*=3, scale=80µm). **G**, cells were treated with 10µM erastin in the presence or absence of 2-8% ascites from liver cirrhosis (LC) or OVCA patients for 24 hours (*n* provided in panel, 24 hours). **H**, micro-organospheres (MOS) were treated with 50µM IKE in the presence or absence of 10-40% OVCA ascites for 72 hours (*n*=5). **I-J**, CAOV3 cells transduced with lentivirus expressing luciferase were resuspended in PBS or human OVCA ascites and IP injected into 6-week-old SCID beige mice. Tumor growth (**I**) was visualized and (**J**) measured as total photon flux (p/s) 10 days after tumor injection via IVIS bioluminescent imaging (*n*=6). Cell viability tests were measured with the CellTiter-Glo® assay. All data represent mean ± s.d. Statistical significance was assessed using correlated-samples one-way or two-way (**F**) ANOVA, and multiple comparisons were adjusted using Holm-Šídák’s method (**D-H**). Statistical significance for **J** was assessed using the two-tailed Student’s *t*-test method.

We then investigated whether ferroptosis protection required the continuous presence of ascites. To test this, we added 8% ascites to CAOV3 cells for 16 hours, after which the ascites was removed, washed with PBS, and the cells were treated with erastin and monitored for cell death for 72 hours (**Fig. 1E**). We used the CellTox^TM^ Green Cytotoxic reagent for monitoring cell death at multiple time points, which emits green fluorescence via binding to DNA of cells with impaired membrane integrity (31). Intriguingly, while CAOV3 cells not previously exposed to ascites died within 24 hours after erastin treatment, CAOV3 cells that had been transiently exposed to ascites or had continuous ascites exposure were still viable after 72 hours of erastin treatment (**Fig. 1F**). These results indicate that transient exposure to ascites induces a ferroptosis-resistant state that persists even after its removal. The results, together with the ability of ascites to protect against ferroptosis triggered by *GPX4* knockdown (**Supplementary Fig. 1C**), also further rule out the possibility that some ascites components inactivate various ferroptosis-inducing agents to protect against ferroptosis.

We then wanted to further test the reproducibility of this ferroptosis protection phenotype in additional ascites samples and its potential relationship to disease settings. Other than advanced OVCA, many patients with liver cirrhosis (LC) also develop ascites (32). Therefore, we compared the ferroptosis protection effects of a dose-titration of 16 OVCA ascites samples and eight LC ascites samples against ferroptosis induced by erastin in CAOV3. While potential variability between individual ascites samples may limit the reliability of direct comparisons, we observed that LC ascites samples, in contrast to OVCA ascites, had less ferroptosis protection as they require a significantly higher dosage and exhibit less consistency in their protective effects (**Fig. 1G**). Next, we confirmed the ability of ascites to rescue against ferroptosis in organoids that had been derived from two OVCA patients with different cancer types – serous ovarian carcinoma (MOS 01), and adenocarcinoma with squamous differentiation of the ovary (MOS 02). Micro-organospheres (MOS) (33) developed from these two patient-derived organoids (PDOs) were treated with ascites and IKE (34), a modified and more potent, soluble form of erastin. Our results show that ascites protected both MOS lines against IKE-induced ferroptosis in a dose-dependent manner (**Fig. 1H**), confirming that ferroptosis protection by ascites can be extended to patient-derived organoids.

Ferroptosis has been proposed as a potential tumor suppression mechanism, and detached tumor cells have been shown to be especially sensitive to ferroptosis (20). Therefore, we wished to determine whether the ferroptosis protection provided by ascites may affect the growth of detached ovarian tumor cells in the peritoneal cavity to model peritoneal tumor growth. To this end, CAOV3 cells transduced with a luciferase reporter and resuspended in either PBS or human ascites were injected intra-peritoneally (IP) into immunodeficient mice. The mice then received cell-free PBS or ascites IP injections 5 and 10 days after tumor injections and were subsequently monitored for tumor growth via IVIS imaging on day 10. We observed that ascites injections significantly increased tumor growth of CAOV3 in the peritoneal cavity (**Fig. 1I-J**). Additionally, compared with PBS injections, tumors harvested from mice with the ascites had lower lipid peroxidation levels, indicative of reduced levels of naturally occurring ferroptosis (**Supplementary Fig. 1K-L**). Together, these results show that OVCA ascites provides a robust, long-lasting, and specific protection against ferroptosis, which may enhance the survival and spread of metastasizing OVCA cells in the peritoneal cavity.

### Ascites lipids dictate ferroptosis protection

Next, we wanted to investigate which components in ascites are responsible for the ferroptosis protection. The acellular components of ascites consist of at least lipids, proteins, and small metabolites (35). To remove different components, we used dialysis to remove small metabolites from ascites, heat to inactivate proteins, and the biphasic separation method (36) to remove the lipids. Triglycerides and cholesterol measuring kits were used to ensure the removal of lipids (**Supplementary Fig. 2A-B**), and protein levels were measured by BCA assay to normalize different samples (**Supplementary Fig. 2C**). When comparing the effects of removing various components from ascites on its ability to protect against ferroptosis, we found that delipidation, but not dialysis or heat inactivation, abolished ascites-mediated protection against erastin-induced ferroptosis in CAOV3 cells (**Fig. 2A**). The importance of lipids in ferroptosis protection was repeated using additional OVCA cell lines (**Supplementary Fig. 2D**), as well as with an additional ferroptosis-inducer, the GPX4 inhibitor, RSL3 (**Supplementary Fig. 2E**). Together, these results show that the lipid components in ascites are the primary factor in protecting against ferroptosis.

**Fig 2.**
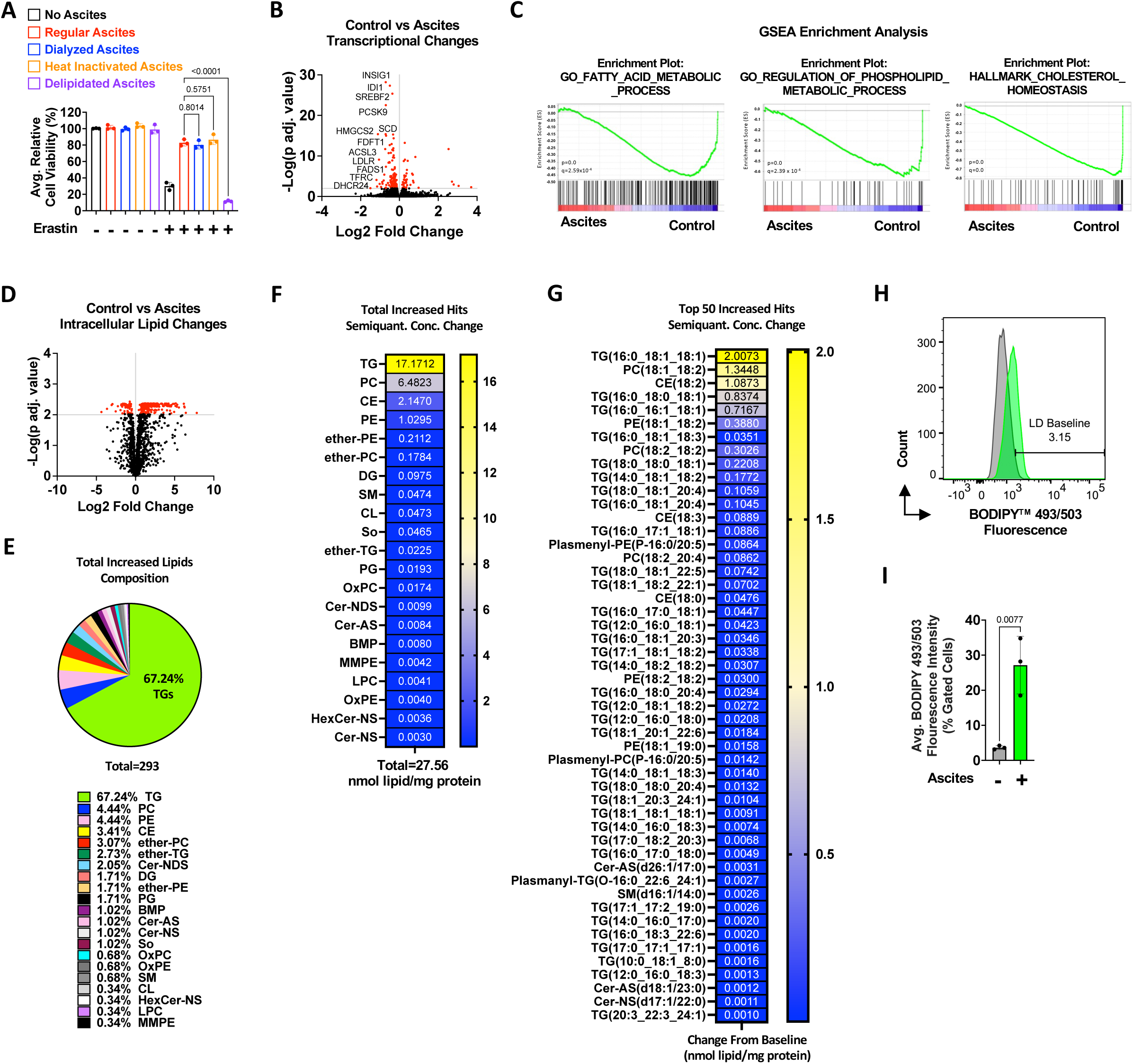
Ascites lipids dictate ferroptosis protection. **A**, cell viability was assessed in CAOV3 cells treated with 2% regular, dialyzed, heat-inactivated, or delipidated OVCA ascites and 10µM erastin (*n*=3, 24 hours). **B**, Volcano plot of the differentially expressed genes (DEGs) of RNA-seq analysis for the transcriptome changes of CAOV3 and TOV21G cells treated with 10% ascites for 16 hours. DEGs from each cell line were combined for a pairwise comparison of total DEGs under ascites-treatment (*n*=3/cell-line, adj. *P*=0.01). **C**, Gene Set Enrichment Analysis (GSEA) for the combined DEGs for the indicated gene-sets from CAOV3 and TOV21G. **D**, structural lipidomic pairwise comparison analysis for CAOV3 cells treated with 5µM erastin with or without 10% ascites for 16 hours (*n*=5, adj. *P*=0.01). **E**, pie-chart of the lipid composition and relative abundance of all significantly increased intracellular lipids in the ascites-treated cells. **F-G**, heat map of all (**F**) significant lipid semiquantitative concentration increases and (**G**) top 50 specific lipid species increases (nmol lipid/mg protein) in erastin and ascites treated cells. **H-I**, CAOV3 cells were treated with 10% ascites and lipid droplet levels were measured via flow cytometry analysis of BODIPY^TM^ 493/503 staining (*n*=3, 16 hours). Cell viability tests were measured with the CellTiter-Glo® assay. All data represent mean ± s.d. Statistical significance was assessed using the two-tailed Student’s *t*-test method (**D**, **I**), and *P* values were adjusted using the Benjamini-Hochberg correction method (**D**).

We then conducted RNA-Seq to identify transcriptional changes of CAOV3 and TOV21G cells exposed to ascites (**Fig. 2B, Supplementary Fig. 2F-G**). Volcano plots of differentially expressed genes (DEG) (**Fig 2B**) and Gene Set Enrichment Analyses (GSEA) (**Fig. 2C**) showed that ascites downregulated many genes in lipid metabolism pathways, further pointing to the relevance of lipid regulation. This prompted us to perform lipidomic profiling of erastin-treated CAOV3 cells upon exposure to ascites. Ascites exposure resulted in a significant increase in intracellular lipids (**Fig. 2D**). Of the 293 lipid species that increased during ascites exposure, 197 (67.24%) were triglycerides (TG) (**Fig. 2E**). In addition to the higher number of TGs, this lipid subclass also showed the most significant increase in total concentration upon ascites exposure (**Fig. 2F-G**). We also performed lipidomic profiling of the ascites sample that was used for cell treatment and found that, of the 293 lipid species increased in ascites-exposed cells, 248 had the same fatty acid distribution as those found in the ascites sample (**Supplementary Fig. 2H**), indicating that the increased lipids originated from the ascites.

Triglycerides are derived from a glycerol backbone conjugated to three fatty acids, either saturated or unsaturated, and they are commonly stored as lipid droplets in the cell (37). They are considered ‘neutral’ lipids that prevent ferroptosis due to sequestering ferroptosis-enhancing polyunsaturated fatty acids (PUFAs) (23). Given the increased amounts of TG content after ascites exposure, we wanted to test whether there was a corresponding increase in the levels of lipid droplets. Thus, we measured lipid droplet content using the BODIPY 493/503 neutral lipid dye with and without ascites exposure. We found that ascites exposure significantly increased lipid droplet levels in CAOV3 cells (**Fig. 2H-I**). Since mono-unsaturated fatty acids (MUFAs) and PUFAs have been reported to have contrasting effects on ferroptosis (30, 38–40), we also analyzed the fatty acid composition of the increased triglyceride to determine whether the ascites-mediated increased TGs are mostly made up of protective MUFAs or ferroptosis-enhancing PUFAs. Our analysis showed the protective MUFA, oleic acid (18:1) to be the most represented fatty acid in the increased TGs (**Supplementary Fig. 2I**). Together, these results identify lipids, specifically triglycerides, as the key components in ascites-mediated protection against ferroptosis.

### Lipid-lowering fibrates mitigate ascites protection against ferroptosis

Given our results, it is likely that the ferroptosis protection of ascites is associated with an increase in protective lipids. Therefore, we tested a small panel of lipid-lowering drugs for their potential to affect ascites-mediated ferroptosis protection (**Supplementary Fig. 3A**). Among the tested drugs, bezafibrate and C-75 Trans mitigated the ascites’ protection against ferroptosis. While we were not able to reproduce the mitigating effects of C-75 Trans, we reproduced bezafibrate’s mitigation of the ascites’ protection against ferroptosis triggered by both erastin (**Fig. 3A**) and RSL3 (**Fig. 3B**). Bezafibrate also mitigated the ferroptosis protection by ascites in multiple types of OVCA cell lines, including TYKNU, TOV21G, and TOV112D (**Supplementary Fig. 3B-D**), as well as when using different ferroptosis inducers, such as IKE and JKE-1674 (**Supplementary Fig. 3E-F**). A similar mitigation of ferroptosis protection was observed using other fibrates, including fenofibrate and ciprofibrate, which are more selective against PPARα (41–43), and a non-fibrate PPARα agonist, GW7647 (**Supplementary Fig. 3G-H**). Importantly, these fibrates did not show notable toxicity on their own but were observed to specifically reverse ascites protection against ferroptosis. Among these fibrates, we found bezafibrate to be more soluble and have the lowest toxicity at baseline (**Supplementary Fig. 3G**). Hence, we focused on bezafibrate in our subsequent experiments.

**Fig 3.**
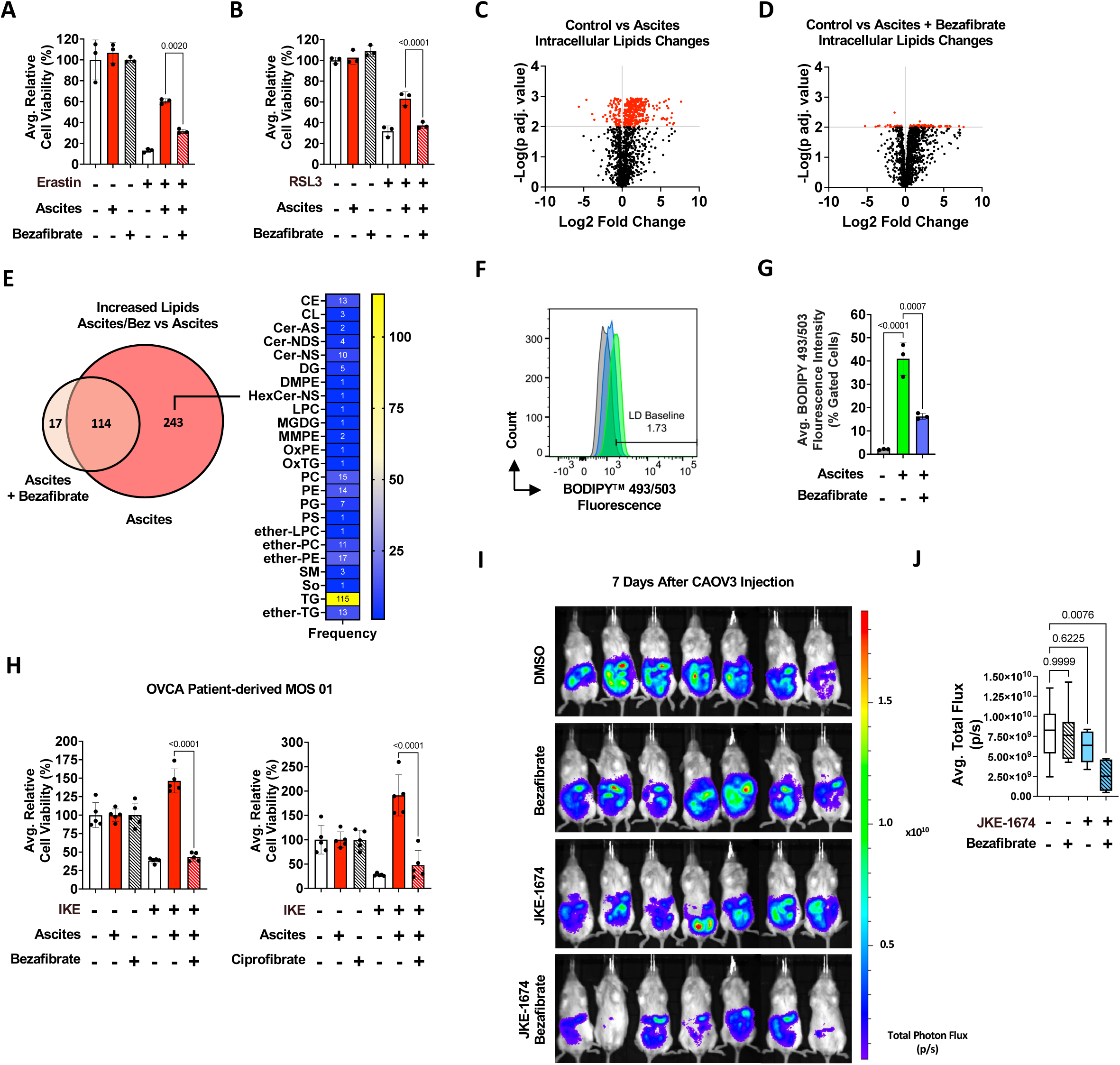
Fibrates mitigate ascites protection against ferroptosis. **A-B**, cell viability of CAOV3 cells was assessed via treatment with (**A**) 10µM erastin or (**B**) 250 nM RSL3, together with 2% ascites, in the presence or absence of 200µM bezafibrate (*n*=3, 48 hours). **C-D**, Volcano plots of the different lipid species from lipidomic comparison analyses for CAOV3 cells treated 10% ascites (**C**) with or (**D**) without 200µM bezafibrate for 16 hours (*n*=5, adj. *P*=0.01). **E**, Venn diagram comparison of lipid changes between ascites-only versus ascites-and-bezafibrate treated cells. The heat map depicts the class composition of the 243 lipids that remain unchanged with the ascites-and-bezafibrate treatment. **F-G**, cells were treated with 10% ascites in the presence or absence of 200µM bezafibrate and lipid droplet levels were measured via flow cytometry analysis of BODIPY^TM^ 493/503 staining (*n*=3, 16 hours). **H**, micro-organospheres (MOS) developed from an OVCA patient were treated with 20% ascites and 50µM IKE, and 1mM bezafibrate or 1mM ciprofibrate to assess resensitivity to ferroptosis (*n*=5, 72 hours). **I-J**, CAOV3 transduced with lentivirus expressing luciferase were treated with either 800µM bezafibrate, 10µM JKE-1674, or both, 5 hours before being injected IP into 6-week-old NSG mice. Tumor growth was then (**I**) visualized and (**J**) measured as total photon flux (p/s) 7 days after injection via IVIS bioluminescent imaging (*n*=7). Cell viability tests were measured with the CellTiter-Glo® assay. All data represent mean ± s.d. Statistical significance for **C-D** was assessed using the two-tailed Student’s *t*-test method (**D**, **I**) and *P* values were adjusted using the Benjamini-Hochberg correction method (**C-D**). All other statistical significance was assessed using correlated-samples one-way ANOVA, and multiple comparisons were adjusted using Holm-Šídák’s method. All statistical tests were two-tailed where applicable.

We then employed lipidomic profiling to determine how bezafibrate affected the ascites exposure-mediated changes in intracellular lipids. Notably, the combination of bezafibrate with ascites significantly subdued intracellular lipid increases when exposed to ascites alone (**Fig. 3C-D**). A comparison analysis of all increased lipids in cells treated with ascites and bezafibrate showed many ascites-increased TGs were unaltered or significantly lowered by simultaneous treatment with bezafibrate (**Fig. 3E**). Consistently, treatment with a combination of bezafibrate and ascites significantly reduced lipid droplet levels compared to treatment with ascites alone (**Fig. 3F-G**). Having established the lipid-lowering effect of the bezafibrate, we then validated that ascites reduced the activity of the PPAR Response Element (PPRE)-driven luciferase reporter, which was increased by combining bezafibrate with ascites (**Supplementary Fig. 3I**). Among all three PPARs, PPARα may be the most relevant as other PPARα agonists also mitigated the ascites rescue effect (**Supplementary Fig. 3G-H**). Additionally, our RNA-seq showed many PPARα target genes were downregulated by ascites (**Supplementary Fig. 3J**). Collectively, these results suggest that the ferroptosis resensitization by bezafibrate is due to a reversal of changes made by ascites exposure, mainly through the PPARα pathway and via the reversal of the intracellular lipid increase in the form of lipid droplets.

We then wanted to confirm the ferroptosis resensitizing effect of bezafibrate in OVCA PDOs. Consistent with our observation in OVCA cell lines, both bezafibrate and ciprofibrate reversed the ascites-mediated protection against the IKE-mediated ferroptosis of OVCA PDOs (**Fig. 3H**). Subsequently, we wanted to determine whether bezafibrate affected the tumor growth of detached tumor cells in the peritoneal cavity in the presence of mouse ascites. We injected immunodeficient mice with CAOV3 cells treated with DMSO vehicle, the ferroptosis-inducer JKE-1674, bezafibrate, or combined JKE-1674 and bezafibrate. We chose JKE-1674 because it is available as an oral administration (34) and requires a lower dosage (**Supplementary Fig. 1C-D**). We monitored peritoneal tumor growth of CAOV3 for seven days via IVIS imaging and observed that, as expected, cells treated only with JKE-1674 showed similar tumor growth as the control, possibly due to protection provided by ascites developed in these mice (**Supplementary Fig. 3K**). However, a combination of JKE-1674 and bezafibrate treatment significantly reduced peritoneal OVCA growth (**Fig. 3I-J**). Taken together, our findings show that using fibrates may mitigate the ferroptosis protection of OVCA cells and PDOs by ascites and reduce the peritoneal spread in OVCA murine models.

### HMGCS2 and TFRC direct ascites-mediated protection against ferroptosis

To elucidate the underlying mechanisms involved in the ferroptosis protection by ascites that were reversed by bezafibrate, we conducted RNA-Seq of CAOV3 cells treated with bezafibrate. As expected, bezafibrate exposure upregulated the expression of many downstream targets of PPARα (**Fig. 4A**). We compiled a list of ascites-regulated genes from RNA-Seq results collected from cells exposed to ascites from two different OVCA patients and compared the ascites-affected gene list to bezafibrate-affected genes. Such comparison analysis identified 13 genes that were repressed by ascites exposure and reversed by bezafibrate (**Fig. 4B**). To prioritize among these 13 genes, we conducted a focused screening and found that ferroptosis was robustly protected by the knockdowns of 3-Hydroxy-3-Methylglutaryl-CoA Synthase 2 (*HMGCS2*) and the transferrin receptor gene (*TFRC*) (**Supplementary Fig. 4A-B**).

**Fig 4.**
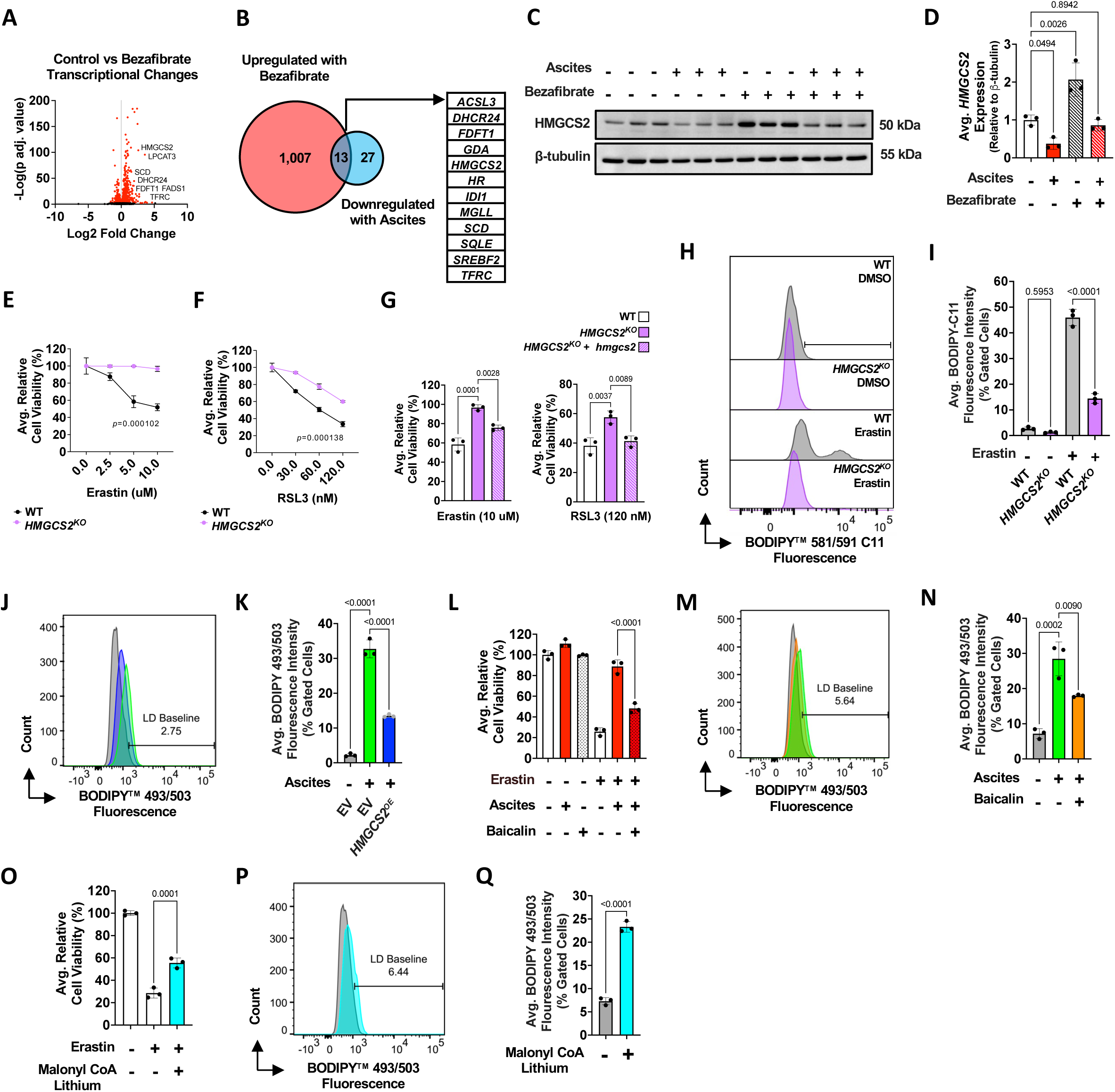
HMGCS2 and TFRC direct ascites-mediated protection against ferroptosis. **A**, Volcano plot of the DEGs of bezafibrate treatment (200µM, 16 hours) from RNA-seq analysis of CAOV3 cells (*n*=3, adj. *P*=0.01). **B**, Venn diagram comparison of a list of oppositely expressed genes from ascites-treated vs. bezafibrate-treated cells. **C-D**, western blot analysis of HMGCS2 protein expression upon 10% ascites treatment from three patients with or without 200µM bezafibrate for 16 hours (*n*=3). **E-F**, ferroptosis sensitivity was compared between CAOV3 wildtype (WT) and *HMGCS2^KO^* cells with (**E**) erastin or (**F**) RSL3 treatment (*n*=3, 24 hours). **G**, HMGCS2 expression was restored in *HMGCS2^KO^* cells with *hmgcs2^OE^* plasmid and ferroptosis sensitivity was compared with 10µM erastin or 125nM RSL3 treatment for 24 hours (*n*=3). **H-I**, lipid peroxidation was measured via flow cytometry analysis of BODIPY^TM^ 581/591 C11 staining in *HMGCS2^KO^* cells that were treated with 5µM erastin for 20 hours (*n*=3). **J-K**, *HMGCS2^OE^* cells were treated with 2% ascites for 16 hours and lipid droplet levels were measured via flow cytometry analysis of BODIPY^TM^ 493/503 staining (*n*=3). **L**, cell viability was assessed via treatment with 10µM erastin in the presence or absence of 20µM baicalin with 2% ascites (*n*=3, 24 hours). **M-N**, cells were treated with 10% ascites in the presence or absence of 25µM baicalin (24 hours) and lipid droplet levels were measured as described (**N**) (*n*=3). **O**, cell viability was assessed in cells treated with 10µM erastin in the presence or absence of 100µM malonyl CoA lithium for 24 hours (*n*=3). **P-Q**, cells were treated with 100µM malonyl CoA lithium (16 hours) and lipid droplet levels were measured as described (**Q**) (*n*=3). Cell viability tests were measured with the CellTiter-Glo® assay. All data represent mean ± s.d. Statistical significance for **D-O** was assessed using correlated-samples one-way ANOVA, and multiple comparisons were adjusted using Holm-Šídák’s method. Student’s *t*-test was performed for **Q**. All statistical tests were two-tailed where applicable.

*HMGCS2* encodes the mitochondrial, rate-limiting ketogenesis enzyme and is a canonical target of the PPARα (44). We observed that its expression was significantly repressed by ascites and reversed by bezafibrate (**Fig. 4C-D**, **Supplementary Fig. 4E**). Importantly, CRISPR-mediated removal of *HMGCS2* also robustly protected against ferroptosis induced by erastin and RSL3 (**Fig. 4E-F**), and ferroptosis sensitivity was restored by expression of the sgRNA-resistant mouse *hmgcs2* (**Fig. 4G**). Moreover, erastin-induced lipid peroxidation was significantly reduced in the *HMGCS2^KO^* CAOV3 when compared with control CAOV3 (**Fig. 4H-I**). In contrast, overexpression of *HMGCS2* enhanced lipid peroxidation levels when compared with empty vector (EV) control in CAOV3 cells (**Supplementary Fig. 4J-K**).

These observations led us to hypothesize that changes in HMGCS2 levels enable the ascites protection against ferroptosis and its subsequent reversal by bezafibrate. Previous studies have shown that HMGCS2 dysregulation can affect lipid accumulation and lipid droplet content (45, 46). Consistently, similar to bezafibrate, the overexpression of *HMGCS2* in CAOV3 mitigated the increase in lipid droplets after ascites exposure (**Fig. 4J-K**).

We then wanted to determine how HMGCS2 modulates lipid droplet content. Given the function of this enzyme in the ketogenesis pathway (44), we first attempted to measure changes in 3-hydroxybutyric acid (BOH) and acetoacetic acid (AcAc) after ascites exposure or HMGCS2 regulation. However, we were unable to detect changes in ketone levels. HMGCS2 is also established in its regulation of fatty acid oxidation (49–51), and its knockdown has been shown to abolish the process (47). Fatty acid oxidation is the process of breaking down fatty acid molecules for acetyl-CoA generation and cellular energy production and, as a result, causes lipid droplet breakdown (48). Given this, we wanted to validate whether fatty acid oxidation could play a role in the modulation of lipid droplet content and ferroptosis sensitivity. To do so, we assessed the effects of baicalin, a fatty acid oxidation activator via the activation of carnitine palmitoyltransferase 1 (CPT1) (49), the first rate-limiting enzyme of fatty acid oxidation (50). We found that baicalin significantly reduced ascites protection (**Fig. 4L**), as well as subdued ascites-mediated lipid droplets’ increase (**Fig. 4M-N**). Reciprocally, the inhibition of CPT1 by the addition of malonyl CoA lithium protected against ferroptosis (**Fig. 4O**) and increased lipid droplet content (**Fig. 4P-Q**), indicating that fatty acid oxidation influences lipid droplet content and ferroptosis.

Collectively, these results suggest that the ascites protection against ferroptosis is mediated via a decrease in HMGCS2, which leads to a reduction in fatty acid oxidation, thereby increasing lipid droplet content and protecting against ferroptosis. The Cancer Genome Atlas (TCGA) pathology data on patient survival also shows that low expression of HMGCS2 is unfavorable not only in ovarian and liver cancer but across all cancers (**Supplementary Fig. 4L**). This may be consistent with the concept that HMGCS2 downregulation may be associated with ferroptosis resistance, tumor progression, and poor clinical outcomes.

### Oleic acid regulation of TFRC influences ascites-mediated protection against ferroptosis

Another ascites-downregulated gene that was reversed with bezafibrate was *TFRC*, which encodes transferrin receptor protein 1 (TfR1) and mediates the cellular iron uptake via transferrin (51). Previous literature shows TFRC is upregulated during ferroptosis (52), and its expression is associated with sensitivity to ferroptosis-inducing agents (53, 54). Consistently, we observed that knockdown of *TFRC* robustly protected against ferroptosis (**Supplementary Fig. 4A**) and confirmed that ascites repressed the mRNA levels of *TFRC* (**Supplementary Fig. 4F**). Importantly, ascites significantly reduced *TFRC* membrane expression and subsequent labile iron increase by RSL3, which were reversed by bezafibrate (**Fig. 5A-C**). These results indicate that, in addition to changes to cellular lipid composition, ascites can also protect against ferroptosis by blocking the increase in functional expression of key ferroptosis-mediators, namely TFRC, which can be reversed by bezafibrate.

**Fig. 5.**
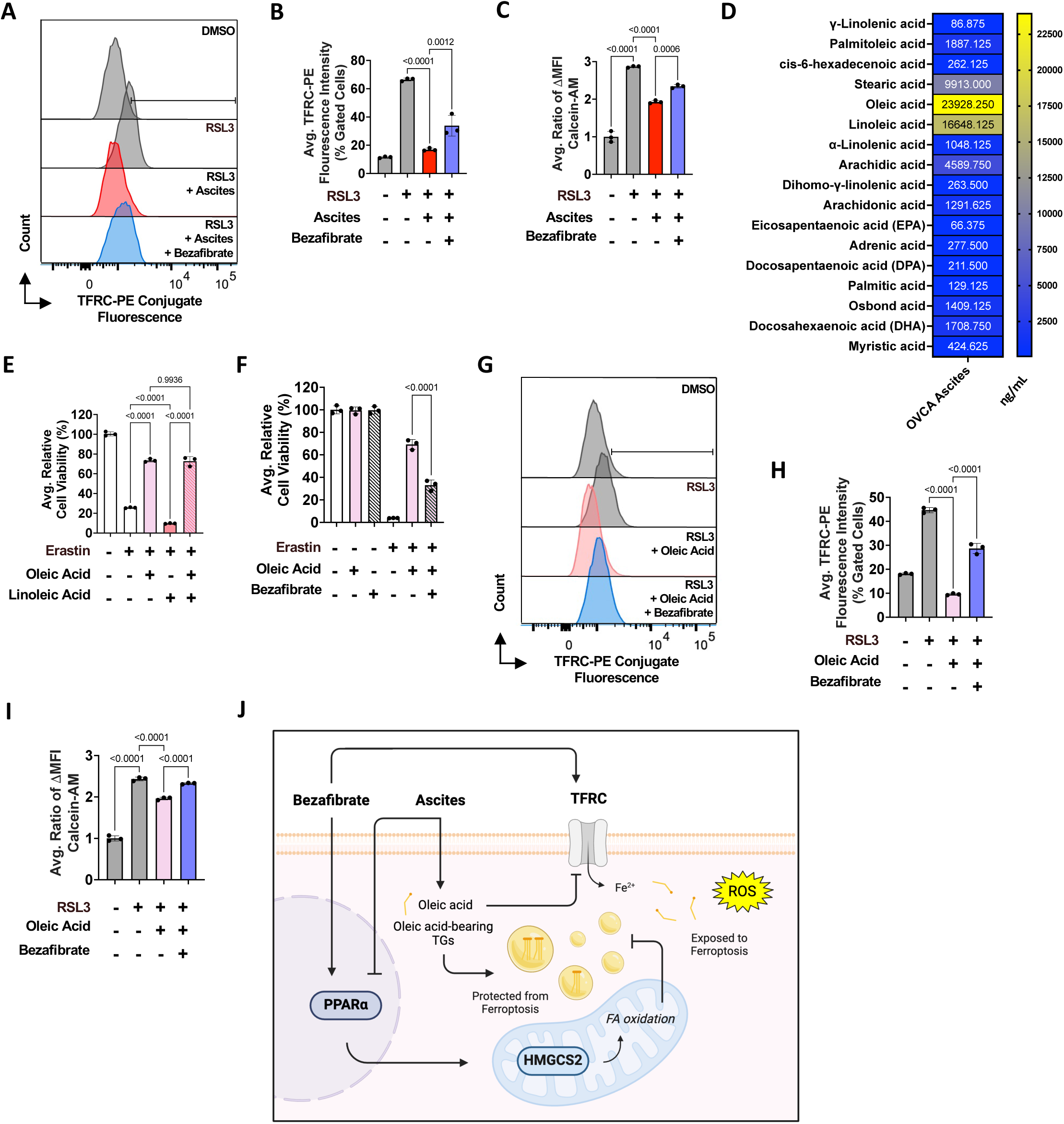
The regulation of TFRC by ascites in protection against ferroptosis. **A-B**, TFRC membrane expression was measured via flow cytometry analysis in cells treated with 2% ascites for 18 hours and 2µM RSL3 and 800µM bezafibrate for 2 hours (*n*=3). **C**, labile iron levels were measured in cells treated with 2% ascites for 18 hours, and 2µM RSL3 and 800µM bezafibrate for 2 hours (*n*=3). Labile iron was measured via calculating the change in Calcein-AM mean fluorescence intensity after treatment with 0.05µM Calcein-AM (15 minutes) and 100µM deferoxamine mesylate (DFO) (1 hour). **D**, heat map of the free fatty acid (FFA) profile in OVCA ascites, measured via FFA lipidomic profiling (*n*=8). **E**, cell viability was assessed in CAOV3 cells that were treated with 10µM erastin in the presence of either 250µM oleic acid, 150µM linoleic acid or both (250µM, 150µM) (*n*=3, 24 hours). **F**, cell viability was assessed in cells treated with 10µM erastin with or without 200µM bezafibrate and 50µM oleic acid for 24 hours (*n*=3). **G-I**, TFRC membrane expression (**G-H**) and labile iron levels (**I**) were measured with using 50µM oleic acid in place of ascites (*n*=3). **J**, mechanism of action model for ascites protection against ferroptosis and its mitigation by bezafibrate. Cell viability tests were measured with the CellTiter-Glo® assay. All data represent mean ± s.d. Statistical significance was assessed using correlated-samples one-way ANOVA, and multiple comparisons were adjusted using Holm-Šídák’s method. All statistical tests were two-tailed where applicable.

We next investigated how ascites regulates TFRC. Previous literature indicates that cis-MUFAs, namely oleic acid, can regulate *TFRC* expression (55). Our structural lipidomic analysis showed oleic acid to be the predominant fatty acid in ascites-mediated cellular TG increases (**Supplementary Fig. 2I**), indicating a predominant presence of oleic acid in ascites and ascites-treated cells. To confirm this, we conducted a free fatty acid analysis of eight OVCA ascites samples and found that oleic acid was indeed the most abundant free fatty acid across all eight samples (**Fig. 5D**). Since we also observed a considerable amount of the PUFA linoleic acid, we added both oleic acid and linoleic acid together in a ratio equal to what was found in OVCA ascites (1.5:1.0) and found such combination still protected against ferroptosis (**Fig. 5E**). Furthermore, bezafibrate had a ferroptosis resensitizing effect on oleic acid-mediated protection, similar to its effect on ascites-mediated protection (**Fig. 5F**). Given the relevance of oleic acid, we further validated that oleic acid does indeed show similar effects to that of ascites on TFRC membrane expression and labile iron levels, reversed by bezafibrate (**Fig. 5G-I**). Concurrently, these data suggest that, in addition to regulating HMGCS2 and intracellular lipids, the decrease in iron via TFRC regulation may contribute to ferroptosis protection by further preventing increased labile iron, which propagates lipid peroxidation.

## Discussion

Our results have revealed an important role of ascites in protecting OVCA cells from ferroptosis, which may enable their growth and spread in the peritoneal cavity. We show that ascites from OVCA patients robustly protects OVCA cells and organoids against ferroptosis, which can be reversed by PPARα agonists. Given the vulnerability of metastatic cancer cells to ferroptosis (56), we found that ascites may protect against such ferroptosis susceptibility to enhance peritoneal growth. Importantly, mitigating such ferroptosis protection by bezafibrate may reduce peritoneal growth when exposed to ferroptosis inducers.

Furthermore, based on the genes showing opposite patterns of expression in ascites-treated versus bezafibrate-treated cells, we report that the protective effects of ascites exposure are mediated via regulating the levels of *HMGCS2* and *TFRC*. *TFRC* regulation is well established in ferroptosis due to its role as an iron importer, which is one of the main propagators of lipid peroxidation (51–54). Hence, its regulatory effects on ferroptosis are not surprising. The effects of HMGCS2 regulation, however, are more intriguing. Previous studies report HMGCS2 overexpression to be associated with a decrease in lipid droplets (47, 48) and an increase in cell death and oxidative stress (57). However, to our knowledge, HMGCS2 has not been linked to ferroptosis sensitivity.

Interestingly, previous studies have shown PPARα activation to have a protective role against ferroptosis, mainly through its influence on lipid metabolism and its activation via the ligand binding of fatty acids (58). However, while PPARα can be activated via ligand binding of ferroptosis-protecting MUFAs, it can also be activated by ferroptosis-enhancing PUFAs (59, 60), and one study shows oleic acid can bind to but not activate PPARα (61). Within the context of our study, we believe that ferroptosis resensitization by PPARα activation only works in the context of ascites exposure and with reversing the effect of the ascites-mediated cellular lipid changes. Treatment with bezafibrate, or PPARα activation alone, could not promote ferroptosis. Aligned with this, bezafibrate-treated cells injected into mice did not significantly affect peritoneal cancer cell growth. Overall, PPARα manipulation seems to be complicated by many context-specific effects on ferroptosis, but its role as a lipid metabolism regulator is undoubtedly important for the cellular response to ferroptosis.

Our study has obvious therapeutic implications. To the best of our knowledge, ascites, which is observed in more than 90% of patients with metastatic OVCA (1, 7), has never been studied in the context of ferroptosis sensitivity, which is a therapeutically relevant Achilles heel for disseminated cancer cells (20, 56). Furthermore, the influence of intracellular lipid changes on ferroptosis sensitivity and its potential reversal via the use of commonly prescribed fibrates may provide a promising direction for further investigation in OVCA therapeutics. We also believe that our findings may have relevance for the peritoneal metastasis of colorectal cancer, as well as pleural, brain, and spine metastases. Finally, our results further emphasize the possible significance of traveling routes as an important determinant of tumor progression and point of potential therapeutic intervention.

## Methods

### Cell culture

The CAOV3, TYKNU, TOV21G, and TOV112D cell lines were a gift from Zhiqing Huang. HEK293T cells were obtained through the ATCC and used for viral packaging. All cell lines were authenticated and regularly tested for mycoplasma via the Duke Cell Culture Facility (CCF). Cells were maintained in a humidified incubator at 37°C with 5% CO_2_. The Dulbecco’s Modified Eagle Medium (DMEM) (Gibco, 11995065) that was supplemented with 10% fetal bovine serum (FBS) (Cytiva, SH30071.03HI-LSF) and 1% streptomycin-penicillin (Gibco, 15140122) was used for all cell lines. All cells were trypsinized using trypsin-EDTA (Gibco, 25200072).

### Ovarian patient-derived organoid establishment, maintenance, and treatment

OVCA tissue samples were collected at the Duke University Hospital through the Duke BioRepository and Precision Pathology Center (BRPC), which is part of the National Cancer Institute’s Cooperative Human Tissue Network. Samples were collected with written informed consent under a Duke Institutional Review Board approved protocol (Pro000089222). Tissue samples were cut into pieces ∼2 mm^3^ with a sterile scalpel and enzymatically digested in 5mL tubes with 4.7mL DMEM F12 media (Gibco, 11320033) and manufacturer-recommended amounts of H, R, and A enzymes in the Tumor Dissociation Kit, human (Miltenyi Biotec, 130-095-929). Samples were digested for 60 minutes at 37°C in the Roto-Therm Plus (Ward’s Science). Cells and tissue fragments were then filtered through a 70µm filter and centrifuged at 500*g* for five minutes. The supernatant was aspirated and 1.25 x 10^5^ cells were plated per 50µL dome of Matrigel (Corning, 356234).

Ovarian patient-derived organoids (PDO) were maintained in ovarian media, which consisted of DMEM F12 media supplemented with the following components: 10mM HEPES, 1X GlutaMax, 100U/mL Penicillin/Streptomycin, 10nM 17-B-Estradiol, 500nM A83-01, 1X B27 without Vitamin A, 50ng/mL EGF, 25ng/mL FGF7, 100ng/mL FGF10, 10nM [Leu15]-Gastrin I, 10ng/mL HGF, 20ng/mL IGF, 1X N2, 1mM N-Acetylcysteine, 10ng/mL Neuregulin I, 10mM Nicotinamide, 100ng/mL Noggin, 100µg/mL Primocin, 10nM Prostaglandin E2, 100ng/mL R-Spondin 1, 3µM SB202190, and 10µM SB203580 (p38i). All PDO were maintained in a 37°C humidified incubator at 5% CO_2_.

Once the PDO were confluent, media was aspirated, and 1mL of PBS was added to each well to detach the Matrigel domes. The solution was centrifuged at 500*g* for five minutes. 1mL of TrypLE Express (Gibco, 12604013) was used to dissolve Matrigel and dissociate organoids. These mixtures were incubated for five minutes at 37°C and TrypLE was neutralized by adding 5mL of DMEM F12 media with 10% FBS and 1% penicillin/streptomycin. After centrifuging at 500*g* for five minutes, supernatants were aspirated. PDO cell suspensions were used to make MicroOrganoSpheres (MOS) as previously described (33) at a concentration of 50 cells per MOS. MOS were then plated at a concentration of 100 MOS in 50µL of media per well in a 96-well plate and treated with DMSO control, IKE (50µM), and 10, 20, and 40% ascites with or without IKE (50µM). Cell viability was assessed using the Cell Titer-Glo® luminescent cell viability assay kit (Promega, G7570) after 72 hours. Samples that were the most responsive to ascites treatment were subsequently made into MOS and treated with the following conditions: DMSO control, 20% ascites, IKE (50µM), IKE (50µM) and 20% ascites, bezafibrate (1mM), bezafibrate (1mM) and 20% ascites, bezafibrate (1mM) and IKE (50µM), and bezafibrate (1mM) and IKE (50µM) and 20% ascites (*n*=5). The same experimental design was repeated with ciprofibrate (1mM) in place of bezafibrate. MOS plates were imaged daily for three days using an ImageXpress Pico (Molecular Devices) cell imaging system, and cell viability was measured as described above after 72 hours.

### Mouse studies

CAOV3 cells were transduced with a GFP-luciferase reporter (System Bioscience, pGreenFire1-CMV Positive Control, TR011PA-1) and GFP+ cells were sorted via flow cytometry. 0.5 × 10^6^ of FC-sorted cells were resuspended in 100µL PBS or human OVCA ascites and injected intraperitoneally (IP) into 12 6-week-old female immunodeficient mice (Taconic Bioscience, CBSCBG-F). 100µL of cell-free PBS or human OVCA ascites were IP injected 5 and 10 days after initial tumor injections. For the second set of mice experiments, FC-sorted CAOV3 cells were treated with either bezafibrate (800µM), JKE-1674 (10µM), or both for 5 hours. Cells were counted and sorted via a hemocytometer to exclude dead cells using trypan blue, and 2.0 x 10^6^ live cells from each treatment group were resuspended in 100µL PBS and IP injected into 28 6-week-old female immunodeficient mice (Duke DLAR Rodent Breeding Core, NOD.SCID.gamma). Injections were done randomly, but mice were subsequently marked on their tails to keep track of treatment groups. All mice were kept on normal chow and fed ad-libitum. Peritoneal tumor growth was assessed via bioluminescent imaging 7 or 10 days after injections. Mice were euthanized 7 or 10 days after tumor injections by carbon dioxide chamber, and tumors and ascites were collected for molecular analysis. Mice were to be euthanized if they became moribund or met other IACUC defined criteria suggesting pain or distress (e.g., weight loss >15%, ruffled fur, etc.) in the approved protocols by Duke IACUC (Registry Numbers A118-21-06 and A115-22-06).

### Bioluminescent imaging

Peritoneal tumor growth was assessed via bioluminescent imaging. 10 minutes before the imaging, mice were anesthetized with isoflurane (Covetrus, 029405) and injected IP with D-luciferin potassium (MedChemExpress, HY-12591B) dissolved in PBS (150mg/kg). The mice were then imaged using an IVIS Kinetic Imaging System (Caliper Life Sciences). The exposure time was set at 1s. Tumor growth was quantified as the bioluminescence signal (total photon flux) calculated with ‘region of interest’ (ROI) measurement tools in the Living Image software (4.8, Revvity).

### Ascites samples

Human ascites samples from ovarian cancer patients were obtained through the Duke University School of Medicine Ovarian Cancer Research Biobank under protocol ID Pro00013710. Human ascites samples from liver cirrhosis patients were obtained through the BRPC under protocol ID Pro00111040. Upon receipt, samples were centrifuged at 1,000 rpm for 5 minutes and filter sterilized via 0.2μm PES filters. All samples were stored at -80°C. The pathology information of each ascites sample is provided in the Supplementary. Mouse ascites samples were collected via post-mortem paracentesis from 7 to 8-week-old SCID-beige or NSG mice 7-10 days after tumor cell injection. Samples were pooled together and centrifuged at 1,000 rpm for 5 minutes and filter sterilized via 0.2μm PES filters.

### Ascites delipidation, dialysis, and heat inactivation

The ascites delipidation protocol was adapted from a lipid-stripped serum protocol (36). Briefly, 3mL of ascites was stirred with 0.6mL of *n*-butanol (Sigma, 34867) and 2.4mL of di-isopropyl ether (Sigma, 296856) for 30 minutes followed by centrifugation at 4,000*g* at 4°C for 15 minutes. The lower phase was separated using a glass Pasteur pipette (VWR) and stirred with 3mL of di-isopropyl ether for 30 minutes, followed again by centrifugation at 4,000*g* at 4°C for 15 minutes. The lower phase was separated and put in a SpeedVac Concentrator (Savant) and spun at top speed for 1 hour. Then, the sample was dialyzed in a 10,000 MWCO Slide-A-Lyzer^TM^ Dialysis cassette (ThermoFisher Scientific, 66380) in a beaker containing 2L saline solution (9g/L NaCl) for 3 days at 4°C. A control ascites sample of the same volume was added to the beaker, and the saline solution was changed daily. After the dialysis process, triglyceride (Promega, J3160), cholesterol (ThermoFisher Scientific, A12216), and protein (ThermoFisher Scientific, 23225) levels were measured using a FLUOstar Optima plate reader (BMG Labtech). Dialyzed and delipidated protein concentrations were calculated according to a BSA standard curve and normalized to that of regular ascites’ for use in subsequent experiments. For example, if a dialyzed ascites sample was observed to contain only half the protein concentration of its corresponding regular sample, 2x the volume of that dialyzed ascites was used in experiments in comparison to its corresponding regular sample. For heat inactivation of ascites, samples were incubated at 56°C for 30 minutes.

### Cell viability and cytotoxic assays

Cell viability assays were done with the CellTiter-Glo® luminescent cell viability assay (Promega, G7570) following manufacturer’s instructions. 2.5 – 3.0 x 10^3^ cells were seeded in white 96-well plates (Corning, 3903) for the assay. After treatment, 10μl of CellTiter-Glo® reagent was added to each well (containing 100 µl of media), and plates were covered and shaken for 2 minutes and then incubated for 10 minutes. The resulting luminescent signal was then quantified using a FLUOstar Optima plate reader. The CellTox^TM^ Green cytotoxicity assay (Promega, G8741) was used for the visualization and measurement of cell death at multiple timepoints. The CellTox^TM^ reagent was added to the media in a 1:1,000 dilution and the fluorescent signal was read using a FLUOstar Optima plate reader.

### Microscopy imaging

Cells incubated with the CellTox^TM^ Green reagent were imaged using an EVOS FL Imaging System (ThermoFisher Scientific) at hour 0, 24, 48, and 72 of treatment. Pictures were taken at the 10x objective (scale=80μm).

### Constructs and lentivirus viral infections

*Small interfering RNAs (siRNAs)* – Nontargeting siRNA (siNT) was purchased from Qiagen (AllStars Negative Control siRNA, SI03650318). All other siRNAs were purchased from Dharmacon. si*GPX4* (M-011676-01-0005) was used at a 100nM working concentration. si*HMGCS2* (D-010179-03-0002, D010179-04-0002) and si*TFRC* (D-003941-07-0002, D-003941-05-0002) were used at 50nM working concentrations. Silencing was induced using the reverse transfection method. Briefly, siRNAs were diluted in 20µL of Opti-MEM™ I reduced serum medium (Gibco, 11058021) at the indicated concentrations. 0.2µL of Lipofectamine™ RNAiMAX transfection reagent (ThermoFisher Scientific, 13778150) was added, and the resulting mix was incubated for 20 minutes before 2.5 x 10^3^ cells were added to the mix in a 96-well plate. Cells were incubated with the siRNA for at least 24 hours before additional treatments or silencing validations. *Stable and transient overexpression* – DNA purification of all plasmids was conducted with using the Qiagen Plasmid Midi kit (12145). For transient overexpression of *HMGCS2*, 100ng of mouse *hmgcs2* ORF clone with a pCMV6-Entry backbone (Origene, MG208162) was diluted in Opti-MEM™ I (9µL). 0.3µL of the TransIT®-LT1 transfection reagent (Mirus, MIR2305) was added before the mix was incubated for 15 minutes in a 96-well plate. After the incubation time, 2.5 x 10^3^ CAOV3 cells were added to the mix. The cells were incubated with the DNA for at least 24 hours before additional treatments. For generating a stable *HMGCS2* overexpressing cell line, the human *HMGCS2* ORF clone with a pCMV6-Entry backbone (Origene, RC208128) was cut and ligated into lentiviral vector (Origene, PS100069) according to the manufacturer’s guidelines. The successfully ligated construct was validated via restriction digestion, as well as DNA sequencing through the Duke Life Science Facility with a V2 forward primer (5’ AGCAGAGCTCGTTTAGTGAACC 3’). The pLenti-*HMGCS2* and the pLenti-empty vectors were each diluted with the pMD2.G (Addgene, 12259) and psPAX2 (Addgene, 12260) vectors at a ratio of 1:1:0.1 in Opti-MEM™ I and packaged with the TransIT-LT1 transfection reagent with 2 x 10^5^ HEK293T cells. The virus supernatant was collected at 48 and 72 hours post transduction, filtered through a cellulose acetate membrane (0.45µm, VWR, 28145481), and pooled together. 1mL of the collection with 1µg/ml of polybrene transfection reagent (Sigma, TR-1003-G) was added to 1.5 x 10^5^ CAOV3 cells. Transduced CAOV3 cells were selected with 1µg/mL of puromycin (Sigma, P8833) for at least 4 days. For generating cells containing a PPRE reporter, the pGreenFire1-PPRE Lentivector (System Bioscience, TR101PA-P) was transduced in CAOV3 cells using the same transduction method described above. *CRISPR-mediated knockout* - the sgRNA *HMGCS2* lentiCRISPR v2 plasmid was constructed for us by the Duke Functional Genomics Core Facility (sgRNA sequence: sense 5’ GATACTTGGCCAAAGGACGT 3’). CAOV3 cells were transduced with the plasmid using the same transduction method described above.

### Chemicals

*Ferroptosis reagents –* Erastin and JKE-1647 were purchased from the Duke Small Molecule Synthesis Facility. Imidazole Ketone Erastin (IKE) was purchased from MedChemExpress (HY-114481) and RSL3 was purchased from Cayman Chemical (19288). Liproxstatin-1 was purchased from MedChemExpress (HY-12726). *Non-ferroptosis cell death reagents –* Staurosporine (STS) was purchased from ThermoFisher Scientific (328530010). Actinomycin D (AMD) (A1410) and rapamycin (553210) were purchased from Sigma. *Lipid lowering drugs –* Bezafibrate was purchased from Sigma (B7273). Ciprofibrate, fenofibrate, sulfosuccinimidyl oleate sodium (SSO) (HY-112847A), BMS-309403 (HY-101903), C-75 Trans (HY-12364A), MF-438 (HY-15822), and GW7647 (HY-13861) were purchased from MedChemExpress. *Fatty acids –* Oleic acid (90260) and linoleic acid (90150) were purchased from Cayman Chemical. *Fatty acid oxidation reagents* – Baicalin (HY-N0197) and malonyl CoA lithium (HY-136408) were purchased from MedChemExpress. *Iron chelator* – Deferoxamine mesylate (DFO) was purchased from Sigma (D9533).

### Luciferase reporter assay

After treatment, 2.5 x 10^3^ cells were washed 1x with PBS and incubated with 1mM D-luciferin diluted in DMEM media for 5 minutes at 37°C. Luminescence signal was quantified via a Varioskan Lux plate reader (ThermoFisher Scientific).

### Flow cytometry analysis

For lipid peroxidation staining, 1.5 x 10^5^ CAOV3 cells were washed 1x with PBS after treatment and stained with 10µM of BODIPY™ 581/591 C11 (Invitrogen, D3861) diluted in DMEM media at 37°C for 1 hour, followed by 1x PBS wash and resuspension in 300µL of EDTA-Trypsin and 500µL of PBS filtered in flow cytometry tubes (Corning, 352235). Quantifications were done with the BD FACSCanto^TM^ II (Biosciences). For neutral lipid staining, 1.5 x 10^5^ CAOV3 cells were washed 1x with PBS after treatment and stained with 10µM of BODIPY^TM^ 493/503 (MedChemExpress, HY-W090090) diluted in DMEM media at 25°C for 30 minutes, followed by resuspension and quantification as described above. 2% FBS media was used for neutral lipid staining. For antibody staining, after treatment, 2.5 x 10^5^ CAOV3 cells were resuspended in flow cytometry antibody dilution buffer (Cell Signaling, 13616) containing 1:50 dilution of PE conjugated TFRC antibody (Cell Signaling, 82582) and transferred to a 96-well round-bottom plate (Corning, 3799) and shaken at 25°C for 1 hour, followed by 3x PBS washes and resuspension in 500µL PBS in flow cytometry tubes. Quantification was done as described. Calcein-AM (ThermoFisher Scientific, C3099) staining was done as previously described (62, 63). Briefly, after treatment, 2.5 x 10^5^ CAOV3 cells were incubated with 0.05µM of Calcein-AM for 15 minutes at 37°C, followed by 1x PBS wash and incubation with either 100µM DFO or no treatment for 1 hour at 37°C. Cells were then washed again with PBS and trypsinized for flow cytometry. For quantification of labile iron (*F*Δ), mean fluorescence intensity of Calcein-AM and DFO treated cells (*F1*) was subtracted from mean fluoresce intensity of Calcein-AM only treated cells (*F0*). All FC datasets were analyzed via the FlowJo^TM^ 10.10.0 software.

### Western blot analysis

*Cells* – 2.5 x 10^5^ CAOV3 cells were lysed in 200µL of RIPA buffer (Sigma, R02778) containing protease inhibitor (Roche, 04693159001) and vortexed at 1,500 rpm for 30 minutes at 4°C using a MixMate® (Eppendorf). *Tumor tissues* – tumor tissues harvested from the mice were snap-frozen in liquid nitrogen and homogenized using a mortar and pestle with RIPA buffer (300µL/5 mg of tissue) before being vortexed. The lysates were then centrifuged at 16,000*g* for 20 minutes. Protein concentration was determined using the Pierce^TM^ BCA protein assay kit and diluted with the 4x Laemmli sample buffer (BioRad, 1610747) before denaturation at 95°C for 5 minutes. Equal protein amounts (15-20µg) were run on 10-12% SDS-PAGE gels, followed by a semi-dry transfer using a 0.45µm PVDF membrane (ThermoFisher Scientific, 88518). Membranes were blocked with 5% milk blocking buffer for 1 hour at 25°C, probed with primary antibodies diluted in 5% BSA overnight at 4°C, followed by 3 5-minute washes by 1x TBST and probing with horseradish peroxidase conjugated secondary antibodies (Cell Signaling, 7074, 7076) for 1-1.5 hours at 25°C. Membranes were subsequently washed 3x for 15 minutes with 1x TBST before imaging using a ChemiDoc^TM^ Imaging System (BioRad). SuperSignal^TM^ West Pico (ThermoFisher Scientific, 34577) and Femto (ThermoFisher Scientific, 34096) chemiluminescent substrate kits were used for developing the signals. *Primary antibodies* – Malondialdehyde (1:1,000, abcam, ab27642) HMGCS2 (1:1,000, Cell Signaling, 20940), and β-tubulin (1:1,000, Cell Signaling, 2128).

### qRT-PCR analysis

*Cells* – 1.5 x 10^5^ CAOV3 cells were lysed in 350µL of RLT buffer containing 1% β-ME. *Tumor tissues* – tumors harvested from mice were snap-frozen in liquid nitrogen and homogenized using a mortar and pestle with RLT buffer (350µL/20 mg of tissue). The homogenized lysate was then run through QIAshredder columns (Qiagen, 79656). RNA from cells and tissues was extracted using the RNeasy® Mini Kit (Qiagen, 74104) following manufacturer’s guidelines. RNA was reverse transcribed using SuperScript^TM^ IV (Invitrogen, 18090200). qRT–PCR was performed using Power SYBR Green PCR Mix (Applied Biosystems) and a StepOnePlus^TM^ Real-Time PCR System (ThermoFisher Scientific). *Primers* – hCHAC1 – Fwd, 5’ GAACCCTGGTTACCTGGGC 3’, Rev, 5’ CGCAGCAAGTATTCAAGGTTGT 3’; hSLC7A11 – Fwd, 5’ TCCTGCTTTGGCTCCATGAACG 3’, Rev, 5’ AGAGGAGTGTGCTTGCGGACAT 3’; hHMGCS2 – Fwd, 5’ AAGTCTCTGGCTCGCCTGATGT 3’, Rev, 5’ TCCAGGTCCTTGTTGGTGTAGG 3’; hTFRC – Fwd, 5’ ATCGGTTGGTGCCACTGAATGG 3’, Rev, 5’ ACAACAGTGGGCTGGCAGAAAC 3’; hβ-actin – Fwd, 5’ CACCATTGGCAATGAGCGGTTC 3’, Rev, 5’ AGGTCTTTGCGGATGTCCACGT 3’; hGAPDH – Fwd, 5’ GTCTCCTCTGACTTCAACAGCG 3’, Rev, 5’ ACCACCCTGTTGCTGTAGCCAA 3’.

### RNA sequencing and analysis

Total RNA was collected with the RNeasy Mini Kit and samples (*n*=3) were sent to Duke SGT facility for RNA-sequencing using the NovaSeq 6000 Illumina Sequencing System. Data was processed by the Duke Analytics Core and Solvuu Inc. The TrimGalore, STAR RNA-seq, and Trimmomatic toolkits were used for trimming and aligning the data. Gene counts were compiled using the HTSeq and RSEM tools, and normalization and differential expression was carried out using the EdgeR and DESeq2 Bioconductor packages with the R statistical programming environment. The false discovery rate (FDR) was calculated to control for multiple hypothesis testing. GSEA was performed to identify gene ontology terms and pathways associated with altered gene expression for each of the comparisons performed. Raw and processed data files are available on NCBI GEO under accession codes GSE281415 and GSE281433.

### Lipidomic profiling

*Structural lipidomic –* 1.0 x 10^6^ CAOV3 cells were treated with the following conditions for 16 hours (*n*=5): DMSO control, erastin (5µM), ascites (10%), erastin and ascites, bezafibrate (200µM), bezafibrate and ascites. After treatment, cells were washed 3x with PBS, trypsinized, and resuspended in 200µL of PBS. A set of T_0_ samples were also collected before treatment. Cell samples and the corresponding ascites sample used for their treatment were sent to BPGbio Inc. for lipidomic profiling. Semiquantitative concentration (nmol lipid/mg protein) of 1,840 lipids were measured via UHPLC-MS/MS. In brief, lipids were extracted using the Bligh-Dyer method, followed by evaporation and reconstitution in 100% isopropanol. Samples were analyzed on an Agilent 1290 LC system coupled to a Thermo Q-Exactive Plus mass spectrometer, using reversed-phase UHPLC (Waters Acquity Premier CSH C18 column) and data-dependent iterative-exclusion MS/MS. Lipids were identified using the LipidMatch Flow software, and peak areas were normalized using appropriate isotopically-labeled internal standards (Avanti Equisplash) and by protein content measurement using a Pierce^TM^ BCA assay. *Long-chain free fatty acid lipidomic* – Eight OVCA ascites samples were sent to Duke Proteomic and Metabolomic facility for a custom LC-MS/MS profiling of 22 long-chain fatty acids (C12-C24). Samples were mixed with 100% methanol at a ratio of 1:3 and stored at -20°C for 20 minutes, followed by centrifugation at 2,000*g* for 5 minutes at 10°C. Samples were analyzed with the Sciex QTrap 6500+ system (Framingham, MA) with Waters Acquity I-class plus UPLC. The internal standard solution used included the following: d4-stearic acid, d3-arachidic acid, and d17-oleic acid are 50μg/mL; d4-linoleic acid, d8-arachidonic, d6-mead acid, d5-DPA, d5-α-linolenic acid, and d6-dihomo-γ-linolenic acid at 1.25μg/mL. Software Analyst 1.7.1 was used for data acquisition. Data analysis was done with the Skyline software (daily version 22.2.1.278).

### Structural lipidomic analysis

Pairwise differences in lipid abundance between treatment groups was determined with the two-tailed Student’s *t*-test method and calculation of the log2FoldChange. *P* values were FDR-adjusted using the Benjamini-Hochberg method (*n*=5, adj. *P*=0.01). For pie chart analyses, based on pairwise comparison groups, each significantly increased lipid species was compiled with its corresponding lipid class and the sum total and composition percent of each class was calculated and visualized using the Prism 10 software (GraphPad). For calculating semiquantitative concentration change of lipid species or lipid class after ascites exposure, semiquantative values of each significantly increased lipid were subtracted from the value in the untreated control. Heat maps were generated using Prism. For calculating total number of each UFA detected in increased triglycerides after ascites exposure, significantly increased TGs (in comparison to untreated control) were separated, and each UFA present was manually counted and tallied together. For comparing the lipid species of ascites sample versus ascites-treated cells, undetected lipids were filtered out and common list elements were analyzed between ascites sample lipids and significantly increased lipids (in comparison to untreated control) using excel. Other Venn diagrams were created following a similar procedure in excel by comparing common list elements between two groups.

### Schematics

Schematic in Figure 5j was created with BioRender.

### Statistics

*n* represents numbers of biologically independent replicates, which are included in each figure legend and illustrated by individual data points in each panel. Sample size was chosen based on our previous experience and literature reports. Specific information on statistical analyses is included in each figure legend and calculated via the Prism 10 software (GraphPad). All data represent mean ± s.d. Statistical significance for any experiments containing more than two groups was assessed using correlated-samples one-way analysis of variance (ANOVA), and statistical significance for experiments with more than two groups that involved multiple timepoints was assessed using correlated-samples two-way ANOVA. Multiple comparisons were adjusted using Holm-Šídák’s method. Statistical significance for experiments containing two groups was assessed using the Student’s *t*-test method. For pairwise comparison of lipidomic data, *P* values were adjusted using the Benjamini-Hochberg correction method. All statistical tests were two-tailed where applicable.

## Availability of Data and Materials

All data and reagents supporting the findings of this study are available from the authors upon reasonable request.

## Acknowledgements

We would like to thank Tianai Sun and Yang Zhang for their valuable mentorship and training. We are grateful to Shannon Jones McCall and Duke BRPC, supported by P30CA014236, and the National Cancer Institute’s Cooperative Human Tissue Network (CHTN (RRID: SCR_004446)), supported at Duke University by UM1CA239755. We thank Lucas Li at the Duke Proteomic and Metabolomic Facility, as well as Raj Sasidharan at Solvuu Inc. We extend our gratitude to Carole Grenier, Ai Yang, Chih-Yan Tsai, and Chia-An Ho for their contribution in progressing experiments. This research was supported by the Ovarian Cancer Research Alliance, and the U.S. Department of Defense (W81XWH-17-1-0143 to J-TC, W81XWH-15-1-0486 to J-TC, W81XWH-20-1-0907, HT9425-23-1-0498 to J-TC).

## Author contributions

Y.S. and J-T.C. conceived the experiments and wrote the manuscript with assistance from S.K.M., Z.H., and S-Y.C. Y.S. performed the majority of experiments. D.L.D. conducted organoid experiments. Ascites samples were obtained courtesy of Z.H. and A.B. Z.H., S-Y.C., Y.L., C.F., J.J.A-H., J.W., C-C.L., A.A.M, M.A.K., and D.S.H. collaborated in the discussion and progression of experiments.

## Competing interests

The authors declare no competing interests.

---

**Supplementary Fig 1.**
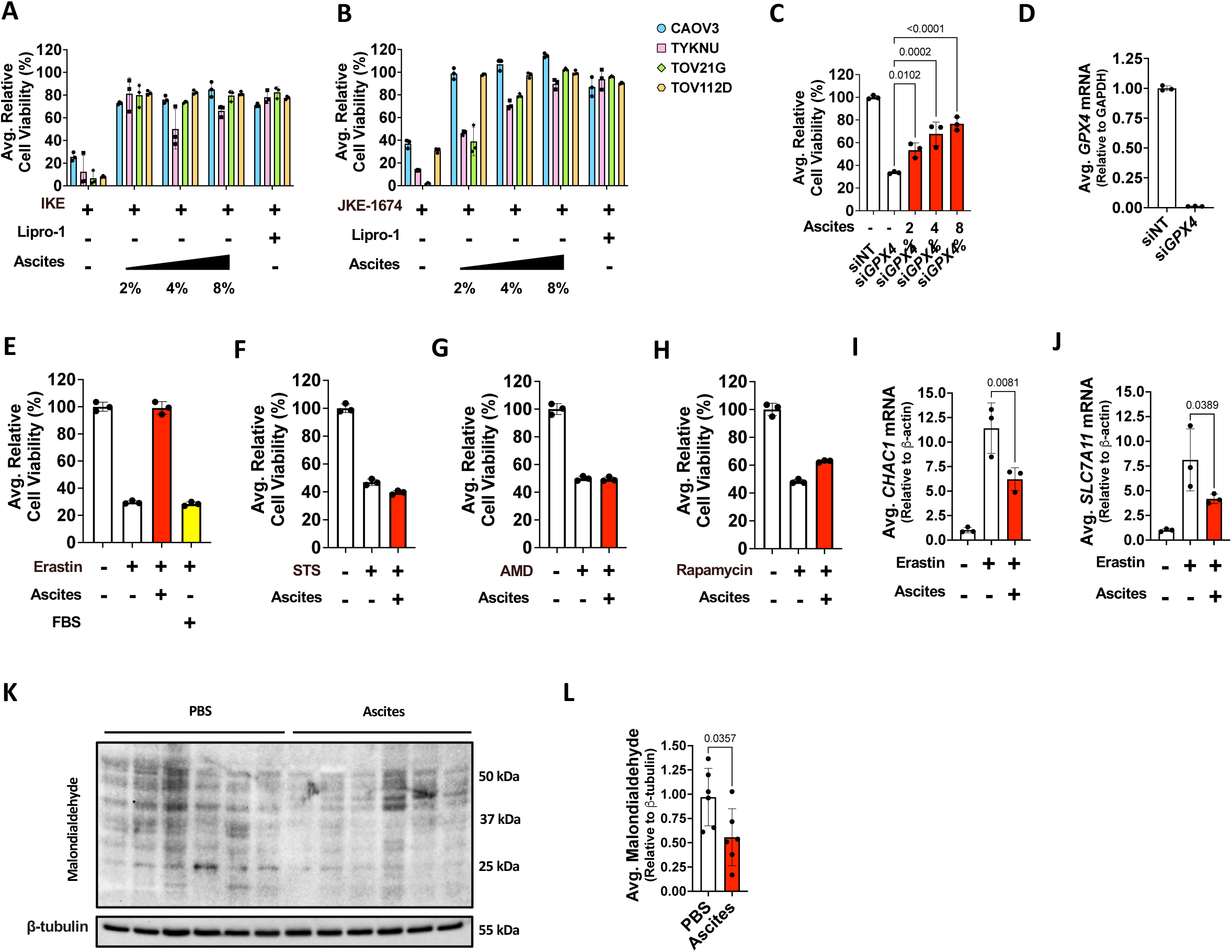
Ascites protects OVCA cells against ferroptosis. **A-B**, CAOV3, TYKNU, TOV21G, and TOV112D cells were treated with (**A**) 5µM IKE or (**B**) 2.5µM JKE-1674 in the presence or absence of 2-8% ascites from three OVCA patients. Liproxtasin-1 (Lipro-1) (2µM) was used as a control (*n*=3/cell line, 24 hours). **C**, validation of ascites rescue against si*GPX4* (100nM) with three different ascites samples in CAOV3 cells (*n*=3, 72 hours). **D**, si*GPX4* knockdown confirmation via qRT-PCR (*n*=3). **E**, CAOV3 cells were treated with 10µM erastin in the presence of either 2% ascites or an additional 2% FBS (*n*=3, 24 hours). **F-H**, CAOV3 cells were treated with (**F**) 200nM staurosporine (STS) for 24 hours, (**G**) 400nM actinomycin D (AMD) for 48 hours, or (**H**) 100nM rapamycin for 48 hours in the presence or absence of 2% ascites (*n*=3). **I-J**, qRT-PCR analysis of (**I**) *CHAC1* and (**J**) *SLC7A11* expression in CAOV3 cells treated with 5µM erastin in the presence or absence of 10% ascites (*n*=3, 16 hours). **K-L**, western blot analysis of lipid peroxidation levels in tumors harvested 10 days after injection of PBS-resuspended and ascites-resuspended CAOV3 cells in SCID beige mice. Signal quantification is depicted in **L** (*n*=6). Cell viability tests were measured with the CellTiter-Glo® assay. All data represent mean ± s.d. Statistical significance for **C**, **I**, and **J** was assessed using one-way ANOVA, and multiple comparisons were adjusted using Holm-Šídák’s method. Statistical significance for **L** was assessed using the two-tailed Student’s *t*-test method.

**Supplementary Fig 2.**
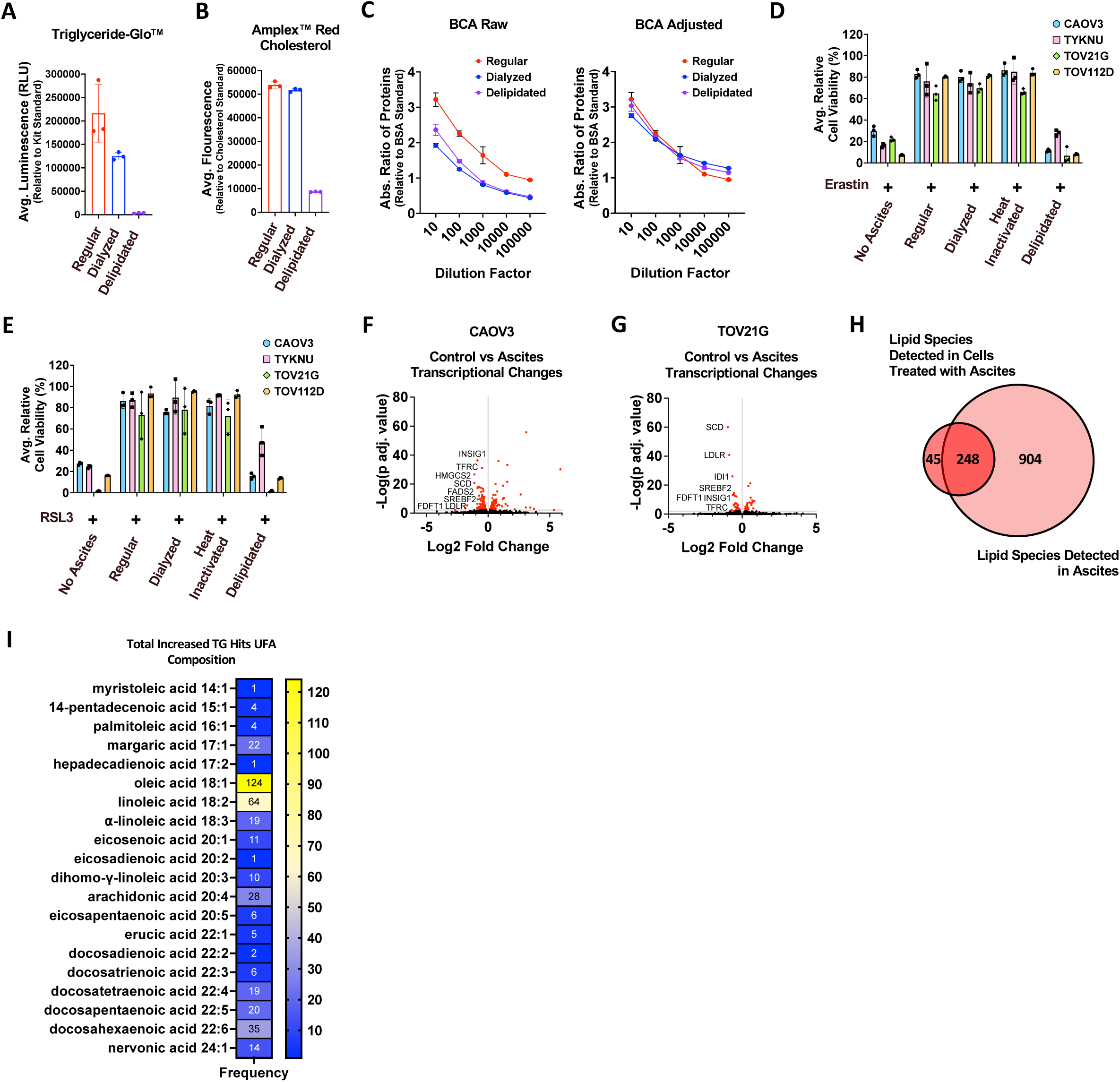
Ascites lipids dictate ferroptosis protection. **A-C**, (**A**) triglyceride, (**B**) cholesterol, and (**C**) protein levels were measured in regular, dialyzed, and delipidated ascites via Triglyceride-Glo^TM^, Amplex^TM^ Red Cholesterol, and Pierce^TM^ BCA assays respectively. Protein levels in dialyzed and delipidated ascites were adjusted using regular ascites as control (*n*=3). **D-E**, CAOV3, TYKNU, TOV21G, and TOV112D cells were treated with 2% regular, dialyzed, heat-inactivated, or delipidated ascites collected from three metastatic OVCA patients in the presence or absence of (**D**) 10µM erastin or (**E**) 250nM RSL3 (*n*=3/cell-line, 24 hours). **F-G**, DEG Volcano plots of the RNA-seq pairwise comparison analyses of (**F**) CAOV3 and (**G**) TOV21G cells that were treated with 10% ascites for 16 hours (*n*=3, adj. *P*=0.01). **H**, Venn diagram comparison of lipid species detected in ascites versus lipid species found to be increased in cells treated with the same ascites (*n*=5). **I**, heat map for sum total of unsaturated fatty acid (UFA) present in increased triglycerides in CAOV3 cells treated with 10% ascites and 5µM erastin for 16 hours. The UFA composition of each triglyceride was recorded and the same UFAs were added together. The sum of each UFA is presented as a number on the heat map (*n*=5). All Cell viability tests were measured with the CellTiter-Glo® assay. All data represent mean ± s.d. Statistical significance was assessed using correlated-samples one-way ANOVA, and multiple comparisons were adjusted using Holm-Šídák’s method. All statistical tests were two-tailed where applicable.

**Supplementary Fig 3.**
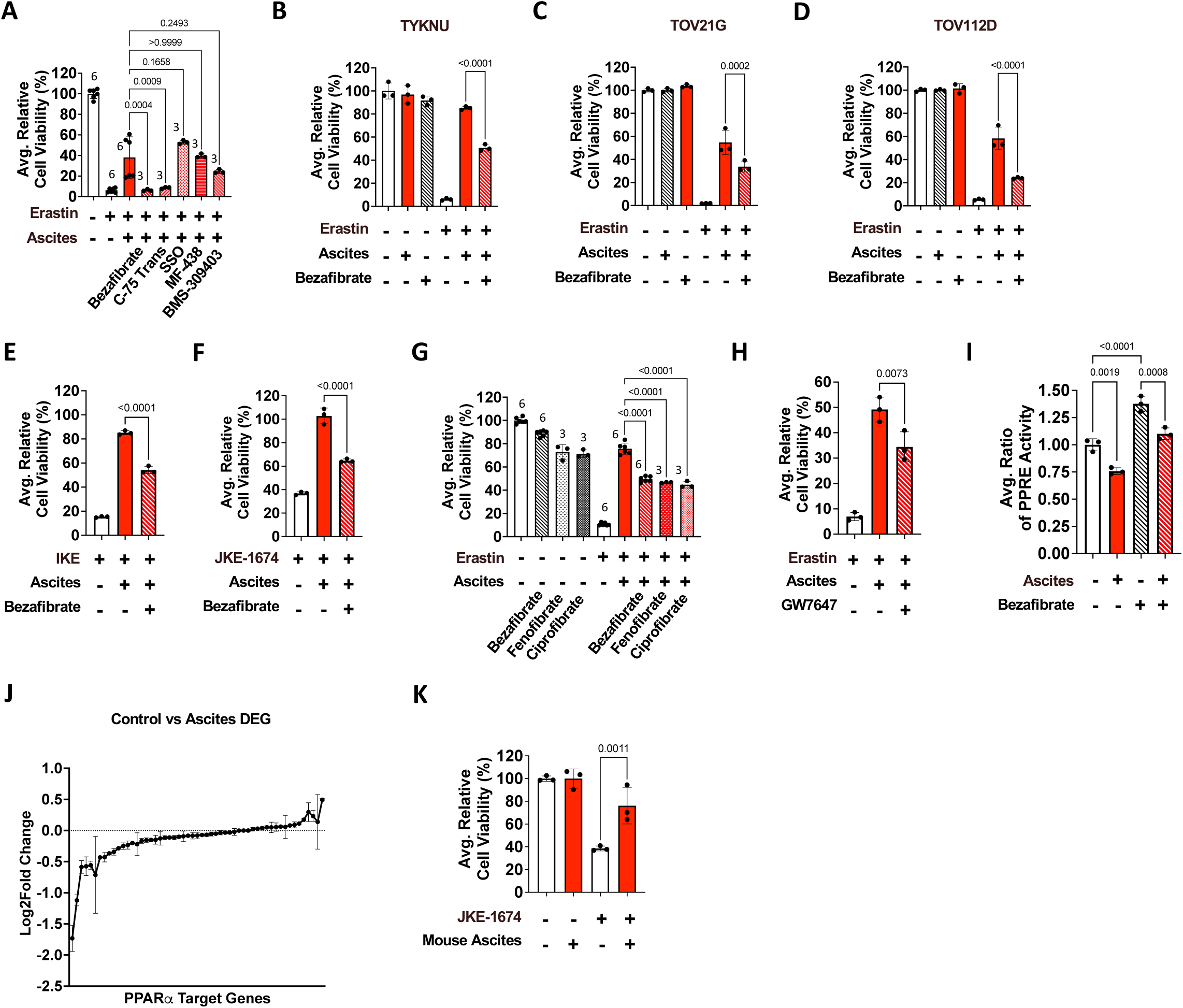
Fibrates mitigate ascites protection against ferroptosis. **A**, Cell viability was assessed in CAOV3 cells treated with 10µM erastin, 2% ascites, and 1mM bezafibrate (PPARα agonist), 0.8mM C-75 Trans (FASN inhibitor), 100µM Sulfo-N-succinimidyl oleate (SSO) (CD36 inhibitor), 50µM MF-438 (SCD1 inhibitor), and 80µM BMS-309403 (FABP4 inhibitor) (*n* provided in panel, 48 hours). **B-D**, (**B**) TYKNU, (**C**) TOV21G, and (**D**) TOV112D cells were treated with 10µM erastin, 2% ascites, and 200µM (**B**, **D**) or 400µM (**C**) bezafibrate (*n*=3, 48 hours). **E-F**, CAOV3 cells were treated with (**E**) 5µM IKE or (**F**) 2.5µM JKE-1674, 2% ascites, and 200µM bezafibrate (*n*=3, 48 hours). **G-H**, CAOV3 cells were treated with (**G**) 10µM erastin, 2% ascites, 200µM bezafibrate, 60µM fenofibrate, 500µM ciprofibrate, or (**H**) 250nM GW7647 (*n* provided in panel, 48 hours). **I**, CAOV3 cells were transduced with a PPRE-luciferase reporter constructed and treated with 2% ascites in the presence or absence of 800µM bezafibrate. Luminescence was measured after 1mM D-luciferin addition (*n*=3, 16 hours). **J**, The log-transformed changes in CAOV3 PPARA target DEGs after 10% ascites exposure (*n*=3, 16 hours, adj. *P*=0.01). The target genes were determined via the PPARGene database (http://www.ppargene.org/index.php), which provides all verified PPARα target genes. **K**, ascites from 7-week-old NSG mice were collected after IVIS imaging. All ascites was pooled together and 8% was added to CAOV3 cells with or without 2.5µM JKE-1674 treatment to assess ferroptosis protection (*n*=3). Cell viability tests were measured with the CellTiter-Glo® assay. All data represent mean ± s.d. Statistical significance was assessed using correlated-samples one-way (ANOVA), and multiple comparisons were adjusted using Holm-Šídák’s method. All statistical tests were two-tailed where applicable.

**Supplementary Fig. 4.**
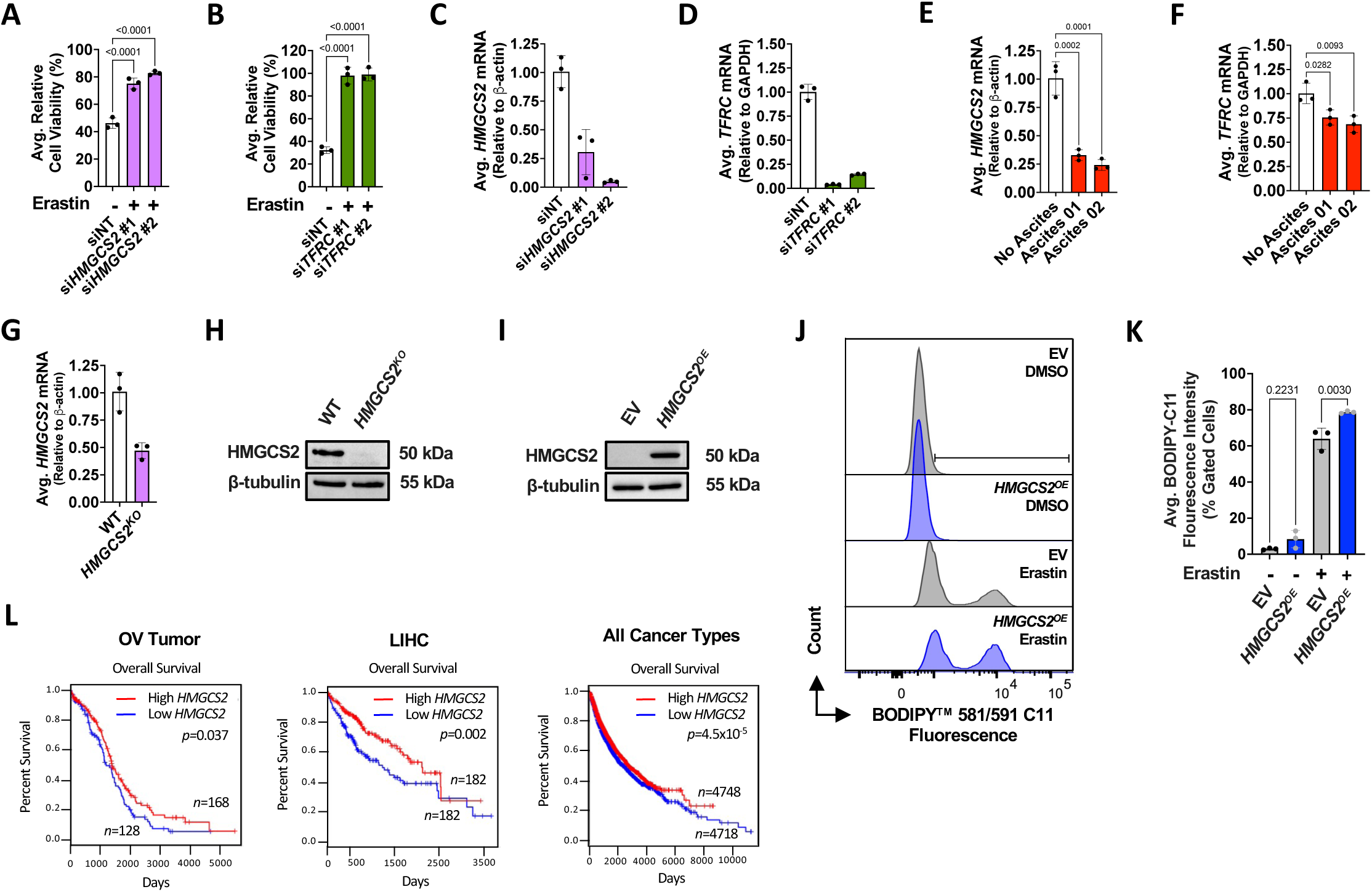
HMGCS2 and TFRC direct ascites-mediated protection against ferroptosis. **A-B**, Cell viability was measured in CAOV3 cells transfected with (**A**) 50 nM si*HMGCS2* or (**B**) 50 nM si*TFRC* and treated with 10µM erastin for 24 hours (*n*=3). **C-D**, (**C**) *HMGCS2* and (**D**) *TFRC* silencing was validated via qRT-PCR (*n*=3). **E-F**, (**E**) *HMGCS2* and (**F**) *TFRC* downregulation upon 10% ascites exposure for 16 hours was validated via qRT-PCR (*n*=3). **G-H**, knockout of HMGCS2 was validated in *HMGCS2^KO^* cells via (**G**) qRT-PCR and (**H**) western blot analysis (*n*=3). **I**, HMGCS2 overexpression was validated via western blot analysis (*n*=3). **J-K**, cells overexpressing *HMGCS2* were treated with 5µM erastin for 20 hours and (**J**) lipid peroxidation was visualized and (**K**) measured with flow cytometry analysis of BODIPY^TM^ 581/591 C11 staining (*n*=3). **L**, patient overall survival data was extracted from the TCGA database and analyzed via the GEPIA 2 survival analysis tool. All cell viability tests were measured with the CellTiter-Glo® assay. All data represent mean ± s.d. Excluding **L**, statistical significance was assessed using correlated-samples one-way ANOVA, and multiple comparisons were adjusted using Holm-Šídák’s method. All statistical tests were two-tailed where applicable.

## Notes

### Competing Interest Statement

The authors have declared no competing interest.

